# Reward Reduces Motor Fatigability by Increasing Movement Vigour

**DOI:** 10.64898/2026.03.24.713707

**Authors:** Jenny Imhof, Caroline Heimhofer, Marc Bächinger, Sarah Nadine Meissner, Richard Ramsey, Nicole Wenderoth

**Affiliations:** Neural Control of Movement Lab, Department of Health Sciences and Technology, ETH Zürich, Zürich, Switzerland; Neuroscience Center Zurich (ZNZ), University of Zurich, ETH Zürich, University and Balgrist Hospital Zürich, Zürich, Switzerland; Brain-Body Regulation Laboratory, Department of Health Sciences and Technology, ETH Zurich, Zürich, Switzerland; Social Brain Sciences Lab, Department of Humanities, Social and Political Sciences, ETH Zürich, Zürich, Switzerland; Future Health Technologies, Singapore-ETH Centre, Campus for Research Excellence and Technological Enterprise (CREATE), Singapore, Singapore

**Keywords:** motor fatigability, motor slowing, pupillometry, upper limbs, reward, EMG

## Abstract

Reward can enhance motor performance. However, its potential to counteract motor fatigability, a reduction in motor performance during sustained movements, remains underinvestigated. This could be particularly relevant in neurological conditions such as multiple sclerosis, where increased motor fatigability is a prominent symptom. One form of motor fatigability is motor slowing, a decline in movement speed over time evoked by fast, repetitive movements. In this study, we investigated whether the possibility to earn reward attenuates motor slowing, and examined associated changes in muscle activity and pupil size, a putative marker of physical effort. Participants performed a wrist tapping task at maximal voluntary speed with or without the possibility of earning a reward. We found that wrist tapping induced motor slowing and that slowing was significantly reduced by reward. Over time, tapping became more costly as indicated by higher muscle activity and coactivation per tap. This was accompanied by a sustained pupil dilation, which could not solely be explained by tapping speed. These findings suggest that, rather than restoring efficient motor control, reward attenuates motor slowing by allowing participants to access a performance reserve and invest more resources into the task, reflected by increased muscle activation per tap and sustained pupil dilation.

## Introduction

Motor fatigability arises during everyday activities that involve prolonged, repetitive movements. It is characterized by a reduction in motor performance over time. Increased motor fatigability, particularly during submaximal movements essential for daily living, has been associated with neurological disorders, such as Parkinson’s disease or multiple sclerosis^1,2^. To investigate the mechanisms underlying motor fatigability, low-force, fast repetitive movements such as finger tapping can be used in healthy individuals^3–5^. In this context, motor fatigability manifests as a gradual decline in movement speed over time, a phenomenon referred to as “motor slowing”^3^. This type of fatigability is predominantly driven by supraspinal mechanisms, as physiological markers of peripheral or muscular fatigue remain largely unchanged^4,6–9^. Fatigability-related motor slowing has been linked to increased blood oxygen level-dependent (BOLD) activity in the primary sensorimotor cortex (SM1), the supplementary motor area (SMA), and the dorsal premotor cortex (PMd)^3^. Dynamic causal modelling (DCM) indicates a reduction in driving input to premotor areas during motor slowing, implying that brain areas beyond cortical motor areas may contribute to motor slowing^5^.

Given the importance of supraspinal mechanisms to motor slowing, the question arises whether these supraspinal mechanisms may be subject to modulation by motivational factors such as reward. While it is well established that reward can enhance motor performance across different species, most studies have focused on effects observed in non-fatigued states^10–17^. In this context, reward-driven improvements in motor performance are thought to be mediated by the optimization of motor control processes^10,11,16,17^ or by the invigoration of movements^15,17–19^. This invigoration can be reflected in faster movement initiation, increased movement speed, increased movement frequency, greater movement amplitude or stronger force production^19,20^. These findings highlight that reward can lead to accepting higher effort-costs for moving faster or more forcefully. However, whether reward can counteract fatigability-related motor slowing remains largely unexplored.

A physiological marker of effort is pupil size, as pupil dilation has been shown to correlate with effort in both mental (for a review see^21^) and physical tasks^22–26^, with larger pupil dilations corresponding to higher levels of effort. Critically, it is thought that pupil size signals the effort invested in the task^21^. However, pupil size is not exclusively related to effort, it has also been linked to bodily states, such as arousal^27^, and several other processes, including reward processing, whereby pupil size is generally larger during high-reward compared to low- or no-reward conditions^28,29^. For instance, during an auditory oddball task, behaviourally-relevant stimuli elicited greater pupil dilations under high-compared to low-reward conditions, while reaction times were shorter^28^. The same pattern was found in a functional magnetic resonance imaging (fMRI) study investigating pupil responses during reward anticipation: higher rewards evoked larger pupil dilations which, in turn, were associated with higher activity in the salience network. It has been argued that the elevated activity in the salience network mediates an increase in arousal to enhance task performance^29^. These findings suggest that reward-related pupil dilation reflects increased arousal and greater effort investment in the task.

Despite extensive evidence that reward improves motor performance, it remains unknown whether reward reduces fatigability-related motor slowing and how pupil size changes during a motor slowing task. Here, we tested whether reward improved motor performance in the fatigued state and whether these improvements were accompanied by initial or sustained pupil dilation, as a potential physiological proxy for increased effort investment. Fatigability was induced by a repetitive wrist-tapping task at maximal voluntary speed which allowed us to quantify tapping speed together with the muscle activity in the wrist flexor and extensor.

## Methods

The research question, hypotheses, planned analyses, sample size and exclusion criteria of this experiment were pre-registered on OSF (https://osf.io/zmpuq) prior to accessing the data. In the present article, we report the preregistered sample size calculation, all data exclusions, all manipulations, and all measures in the study.

### Participants

A total of 26 healthy participants were recruited via online advertisement and via a student mailing list. One participant was excluded from the data analysis due to not understanding the experimental task and was subsequently replaced to meet our predetermined sample size of 25 participants (age: 27 ± 4 y; 13 females). All participants were between 18 to 35 years old, right-handed, free of medication acting on the central nervous system, with no neurological and psychiatric disorders, and with normal or corrected-to-normal vision by contact lenses. Participants were asked to abstain from caffeine intake four hours before testing as caffeine can influence pupil size^30^. Before the study participation, all participants provided written informed consent. The study was conducted with the approval of the research ethics committee of the canton of Zurich (KEK-ZH 2018-01078) and in accordance with the Declaration of Helsinki. Participation was compensated with 20 CHF per hour, and participants could additionally gain a financial reward of up to 9 CHF.

### Sample size estimation

Prior to conducting the study, we determined our target sample size using two complementary power analysis approaches based on our pilot data (n = 7). Pilot participants completed the same tapping task used in the main study, performing 40s tapping trials, in which a cue appeared after 20s indicating whether the current trial was a neutral (no reward) trial or a reward trial with the possibility to win 1 CHF by tapping faster than in the previous reward trail (see *Behavioural paradigm* for details). For the analysis, the tapping data was aggregated into 10s time bins. First, we used G*Power to conduct a traditional power analysis focusing on the key comparison of tapping frequency between reward and neutral conditions during cue presentation. Our pilot data showed a standardized mean difference of Cohen’s d_z_ = 0.7 for this comparison, and G*Power indicated that n = 24 would provide 95% power to detect this effect size in a one-tailed test (α = 0.05). Second, we conducted a Bayesian power analysis using data simulation to account for the multilevel structure of our design. We simulated 1000 datasets based on a Bayesian multilevel model fitted to our pilot tapping frequency data. The design reflected the experimental design of our main experiment, with a 4 × 2 factorial including the factors *Time* (0–10s, 10–20s, 20–30s, 30–40s) and *Condition* (neutral vs. reward). Our primary interest was the *Time × Condition* interaction, because we expected reward effects to emerge specifically when cues were present. With n = 25, 88% of simulated datasets showed 95% quantile intervals for the interaction effect above zero, indicating robust evidence for the expected effect. Combining both approaches, we selected n = 25 as our target sample size. Full details of both power analyses, including complete G*Power parameters and simulation code, are available in our preregistration (https://osf.io/zmpuq/).

### Behavioural paradigm

Participants were asked to perform a motor slowing task combined with reward manipulation. Therefore, participants had to tap their right wrist horizontally by extending and flexing their wrist as quickly as possible between two force sensors for 40s (taping phase; Figure 1a). After the first 20s of tapping, a visual cue (∪ or ∩) appeared on the computer screen positioned in front of the participant (Figure 1b). The cue indicated whether the current trial was a neutral or a reward trial. Cue assignment was counterbalanced across participants: For all participants with an odd participant number, ∪ indicated the reward and ∩ the neutral condition. For participants with an even participant number, the assignment was reversed. During reward trials, participants had the possibility to win 1 CHF if their average tapping speed over the entire trial (mean of 40s tapping) was faster than during the previous reward trial. During neutral trials, there was no reward. Feedback on the reward earned based on the participants’ performance was provided only at the very end of the experiment. Each tapping phase was preceded by a 5s baseline measurement, during which participants were asked to silently count backwards from 100 in steps of 4. Each tapping phase was then followed by a 35s break. The experiment consisted of 20 trials in total, with 10 trials per condition (neutral vs. reward). The trials were presented in a pseudorandomized manner, so that each condition needed to be presented once, before another condition was presented for a 2^nd^ time.

**Figure 1.**
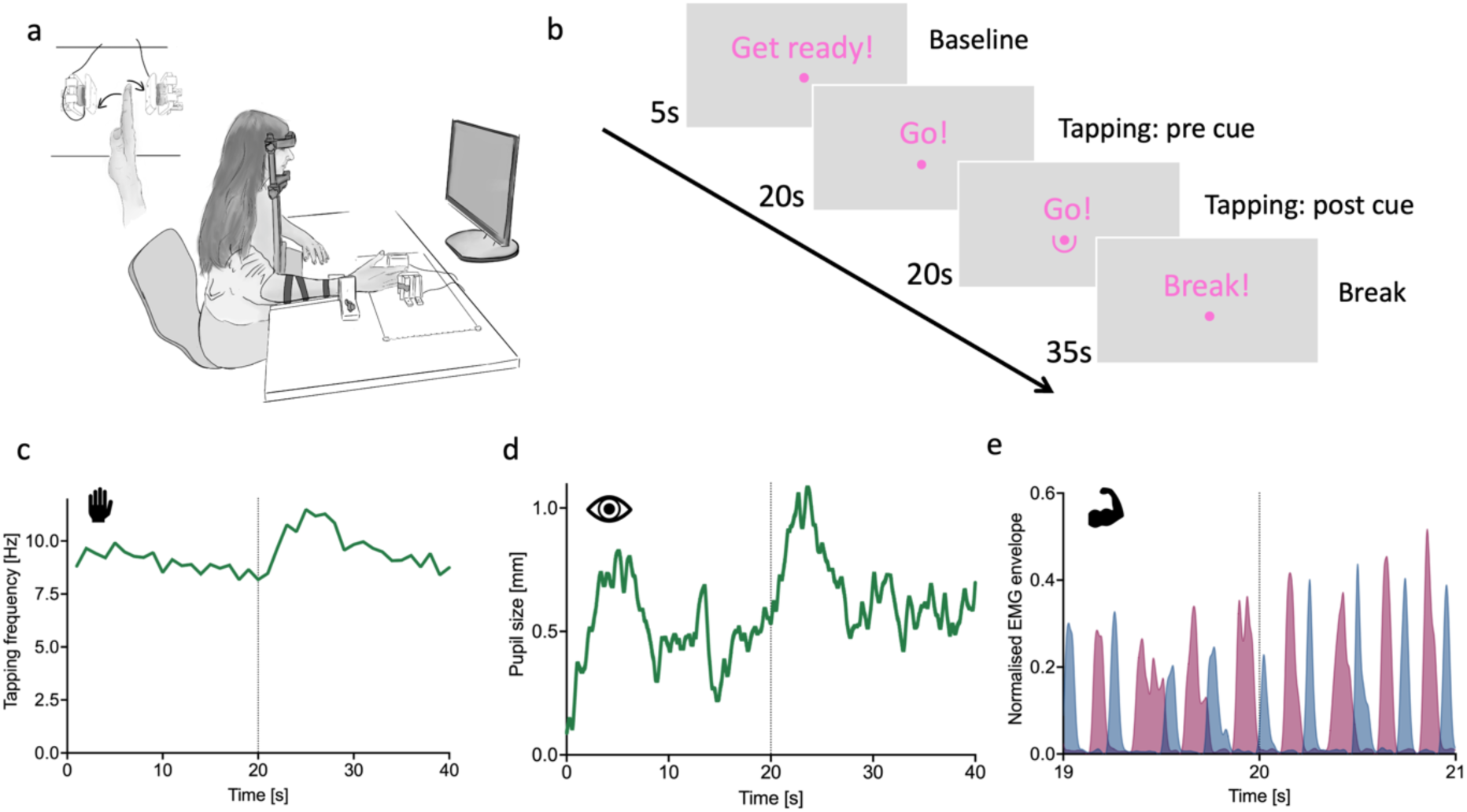
Experimental paradigm. (a) Experiment set up. Participants were asked to tap as quickly as possible between two force sensors, so that they touched the sensors with their fingertips by alternating wrist flexion and extension. To reduce compensatory movements, the participants’ lower arm was fixated. (b) Visual display presented to participants. Each trial consisted of a 5s baseline period indicated by the instruction “Get ready”, a 40s tapping phase signalled by “Go” with a cue presented (∪ or ∩) after the first 20s of the tapping phase, and a break (35s). Example of a reward trial: (c) Tapping frequency in 1-second bins over time, (d) baseline-corrected pupil size time series, (e) electromyographic (EMG) activity showing a 2-second window before and after cue onset. EMG envelopes for flexor carpi radialis (blue) and extensor carpi radialis longus (red) muscles.

### Experimental setup

The experiment was performed in a room with constant dim ambient light to prevent luminance-driven pupil responses and to allow pupil size to vary in both directions (dilation and constriction). Throughout the experiment, participants sat on a stable chair with their head resting against a chin and forehead rest to ensure a stable head position (Figure 1a). The participant’s hand was placed between the two force sensors, with their forearm and elbow supported by the armrest. The forearm was fixated with Velcro to minimize compensatory movements. Additionally, participants were instructed to focus on a fixation dot in the centre of the screen. Taps were recorded using two force sensors (FSR Model 406, Interlink Electronics, Inc., California, USA). Pupil diameter and eye gaze of both eyes were sampled at 60 Hz using the Tobii Pro Nano eye tracker (Tobii Technology) implemented in MATLAB R2023a using the Titta toolbox^31^ in combination with the Tobii Pro SDK (version 1.10.1.2). Before the start of the experiment, the eye tracker was calibrated using a 9-point calibration. To ensure that the changes in pupil size were not driven by changes in lighting, all colours were isoluminant to the grey background (RGB (150 150 150)). Isoluminance was achieved by calculating the relative luminance as a linear combination of red, green and blue components using the following formula: Y = 0.2126 R + 0.7152 G + 0. 0722 B. The formula is based on the principle that green light contributes the most to perceived luminance, while blue light contributes the least (https://www.w3.org/Graphics/Color/sRGB). The experiment was coded in MATLAB 2023a and the stimuli presentation was controlled using the MATLAB-based software Psychtoolbox 3.0.19^32–34^. Besides pupil and tapping data, we also collected the electromyographic (EMG) signal with surface electrodes of the musculus (M.) flexor carpi radialis (FCR) and the M. extensor carpi radialis longus (ECR), the main drivers of the flexion and extension of the wrist, respectively. Using the Bagnoli™ Desktop EMG system (Delsys, Natick, MA, USA), EMG signals were digitally sampled at 1000 Hz through a computer-controlled data acquisition device (CED Power 1401, Cambridge Electronic Design, Cambridge, UK). The signals were amplified and band-pass filtered between 20 and 450 Hz. Both EMG activity and an analogue condition trigger from MATLAB were recorded in Signal software (Cambridge Electronic Design, Cambridge, UK) for later analysis.

### Data analysis

#### Behavioural data pre-processing

The force data recorded by the two force sensors was processed to identify individual taps and to calculate the tapping frequency (Hz). First, for each force sensor, a detection threshold was determined by calculating the root mean square (RMS) of the raw force signal during the tapping phase. This RMS value served as the minimum force threshold to distinguish actual taps from noise and crosstalk between sensors. Only force values exceeding this threshold were considered valid tap signals. Consecutive blocks of above-threshold samples were labelled as individual tap events. Tap onset times were identified by detecting transitions from invalid to valid samples, corresponding to the moment when the applied force first exceeded the RMS threshold. To ensure that each tap represented a distinct force application event, taps occurring within 50ms of each other on the same sensor were filtered out. After processing both sensors independently, the tap times from both sensors were combined and sorted chronologically. Based on the start times of the taps, the tapping frequency was calculated as the inverse of the inter-tapping-interval (ITI), defined as the time between two consecutive force events. Average movement speed for each participant was calculated for both the neutral and reward conditions. Since participants occasionally did not properly hit the target, we implemented a temporal interpolation procedure. Missed taps were inferred when two consecutive events were registered by the same force sensor. In such cases, the ITI was compared to the average time the participant needed to perform a valid movement cycle (defined as the time taken for the first effector to complete the entire movement sequence and return to its starting point). If the ITI fell within one standard deviation of the mean cycle duration (mean ± 1 SD), a single missed tap was assumed on the contralateral side. Interpolated taps were placed at the midpoint between the two detected events using linear interpolation and assigned to the alternate sensor. Intervals exceeding the plausible range (i.e., > mean + 1 SD or < mean – 1 SD) were classified as artifacts. In these cases, the earlier tap registered by the force sensor was retained as the last valid event, and the consecutive tap recorded by the same sensor was removed from further analysis. This procedure preserved the expected alternating tap structure while minimizing distortions in cycle duration estimates. Interpolation rates were 15.7% ± 7.5% for the neutral condition and 15.4% ± 6.7% for the reward condition with no significant differences between conditions (*t*(24) = 0.383; *p* = 0.705).

#### EMG data pre-processing

The EMG signals from the FCR and ECR muscles were processed using the following procedure. First, the EMG signals were high-pass filtered at 20 Hz to remove low-frequency noise and motion artifacts^35^. Second, the signals were rectified and then low-pass filtered at 20 Hz to extract the signal envelopes, providing a smoothed representation of muscle activity over time. The applied low-pass filter cut-off frequency deviates from our preregistered analysis plan, which specified a low-pass cut-off frequency of 10 Hz. However, data inspection revealed that such a setting would have resulted in the loss of meaningful signal components. The resulting signal envelopes of the FCR and the ECR were then normalized to the global peak activity observed in each muscle and were then used to calculate the coactivation index, which was defined as the joint area under the curve (overlap) of the two signals divided by the number of data points^36^. In our preregistration, we specified normalization of the coactivation index to each muscle’s activity, respectively. However, since neither the FCR nor ECR acted as a clear primary mover, we calculated a combined coactivation index. Both the AUCs and the coactivation index were then normalized to tapping speed to account for differences in movement rate across conditions and to isolate differences in muscle activation patterns not solely attributable to speed. In addition, we applied a non-preregistered normalization step by dividing the AUC by the AUC of the first time bin (0-10s). The same normalization was done with the coactivation index to account for differences in outcome measures across conditions in the first two time bins (0-10s 10-20s), likely reflecting insufficient rest periods after reward trials rather than true condition differences (for further details on the order effect, see Supplementary Figure 1). The non-normalized results of the AUC and coactivation index are reported in Supplementary Table 5.

Due to technical issues, EMG data were available for only 6 to 7 trials per condition for all but one participant, with the last 3 to 4 trials missing from the intended 10 trials per condition.

#### Pupil data pre-processing

First, pupil (in mm) and gaze data were visually inspected to ensure that participants focused on the fixation dot throughout the experiment. To avoid bias in pupil size measurements, trials containing large eye movements or saccades (i.e., deviations >∼16° and ∼10° of visual angle on the x and y axes, respectively) were excluded. Next, pupil data was pre-processed following published guidelines and a standardized open-source pipeline^37^. Thus, invalid pupil diameter samples, such as dilation speed outliers and large deviations from the trend line pupil size, were removed using a median absolute deviation (MAD; with the multiplier set to 12 or 30 respectively). Temporally isolated samples with a maximum width of 50 ms that border a gap larger than 100 ms were removed. The remaining valid samples of the right and left eyes were used to calculate the mean pupil size time series. For time points where valid data was only available for one eye, the mean was estimated by applying the dynamic offset between the two eyes, calculated from time points with valid samples for both eyes. The resulting time series was then interpolated to 1000 Hz and smoothed using a low-pass filter with a cut-off frequency of 4 Hz. Data was not interpolated in case the gaps were larger than 250ms. All trials were inspected, and only trials with less than 50% missing data during baseline and tapping phase were included in the analysis. In total, 8 out of 500 trials were excluded from the analysis of pupil data during tapping, with no more than two exclusions per participant. We also analysed pupil data during the subsequent break when participants were resting and excluded 10 additional trials. Finally, to improve statistical power by considering random fluctuations in pupil size over time^38^, subtractive baseline correction was applied for each trial. Therefore, the mean pupil size of the last second of the baseline phase before start of the tapping phase was calculated and then subtracted from every data point of the tapping phase. To confirm that the differences observed in the baseline-corrected pupil time series were not driven by differences in baseline pupil size, the uncorrected mean pupil size of the last second before the onset of the tapping phase was averaged per subject per condition and used in a control analysis.

As one of our statistical analyses aimed to identify specific time points at which the pupil time series differ between conditions, a continuous signal was required. Therefore, baseline-corrected time series were subjected to a cubic spline interpolation at the trial level before averaging across subjects per condition. Cubic spline interpolation was chosen because it better captures the natural variations in pupil size compared to linear interpolation^39^.

### Statistical analysis

We preregistered our primary confirmatory analyses, and deviations from the preregistered plan are detailed in the following section. All statistical analyses were conducted in R Studio (Version 4.4.0), unless otherwise specified. The statistical significance level was set at α = 0.05. Tapping frequency, EMG, and pupil size data were binned into 10s intervals (0–10s, 10–20s, 20–30s, 30–40s) and averaged across trials before statistical testing. Linear mixed-effects models (LMEM) were fitted to the binned data using the lme4 package^40^. For the modelling, we adhered to the “keep it maximal” approach regarding random/varying effects according to Barr et al.^41^. When convergence or singularity issues occurred, model complexity was reduced in a stepwise fashion. For each model, we assessed the distribution of residuals by comparing observed values to the expected uniform distribution using the DHARMa package^42^. Even though LMEM are quite robust, in cases where deviations from normality were detected (Kolmogorov-Smirnov test with *p* < 0.05), we employed a bootstrap approach with 1000 case resamples using the lmeresampler package^43^ to obtain robust confidence intervals and ensure valid statistical inference without relying on distributional assumptions. Since bootstrap confidence intervals were similar to model-based confidence intervals and supported the model findings, detailed comparisons are reported in the supplementary material (Supplementary Tables 1-4). Type III ANOVA tables were generated for each model using the lmerTest package^44^. Partial eta squared (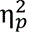) was computed as an effect size for significant factors with the effectsize package^45^. Post-hoc comparisons, adjusted for family-wise error rate (Bonferroni correction), were conducted with the emmeans package^46^.

#### Behavioural analysis

To investigate whether wrist tapping at maximal voluntary speed induced motor slowing and whether participants were already in a fatigued state prior to cue presentation, we first examined changes in tapping frequency over time. Specifically, we were interested in the effect of *Time* within the neutral condition to test that motor slowing occurred. For the reward condition, we focused on the two time bins before the reward cue presentation. Our primary interest, however, was to determine whether the possibility of winning a reward affected motor slowing. The effect of reward was assessed via the interaction of time and condition. We tested these effects by fitting a LMEM with the fixed factors *Time* (i.e., 0-10s, 10-20s, 20-30s, 30-40s), *Condition* (i.e., Reward, Neutral), and their interaction (*Time x Condition)* to the binned tapping frequency data. The model initially included a random intercept for participant and random slopes for the average effects of *Condition*, *Time*, and their interaction. However, due to convergence/singularity issues, we reduced the random effect structure to a random intercept for participant and a random slope for *Condition* only.

#### EMG analysis

To explore the effect of motor slowing and the effect of reward on muscle activity, we applied the same LMEM approach to the coactivation index (as preregistered), as well as to the AUC of the EMG envelopes of the FCR and ECR muscles (the latter was not specifically preregistered) as described for the *Behavioural analysis*. As with the tapping frequency model, we had to stepwise reduce the random effect structure. For the FCR and ECR AUC models, this reduction resulted in a random intercept for participant only. For the coactivation index model, we retained a random intercept for participant and a random slope for *Condition*.

#### Pupil analysis

To assess how pupil size changes during a motor slowing task and whether the effect of reward was reflected in pupil size, the same statistical approach as described for the *Behavioural analysis* and *EMG analysis* was applied to baseline-corrected, binned pupil size data during the tapping phase. In addition to this preregistered analysis, we added tapping frequency as a covariate to the LMEM to explore whether pupil size contains information beyond tapping speed. Both models required stepwise simplification of the random effect structure, resulting in a random intercept for participant and a random slope for *Condition*.

Given that temporal binning reduces the continuous pupil signal to discrete intervals, we conducted a preregistered time-series analysis to test for condition differences across the full temporal trajectory. Baseline-corrected, pupil time series were subjected to a non-parametric paired t-test equivalent implemented in the MATLAB-based SPM1D toolbox for one-dimensional data (SPM1D M.0.4.8; https://spm1d.org/)^47,48^, as the assumption of normality was violated. SPM1D relies on random field theory to draw statistical conclusions at the continuum level of 1D measurements. It assesses the likelihood that smooth, randomly generated 1D sequences would yield a test statistic sequence with a maximum surpassing a specific threshold. While originally developed for the analysis of 1D kinematic, biomechanical, and force trajectories, SPM1D has also been applied to pupil size time-series data^49^.

We pre-registered an additional time series analysis focusing on the cue-evoked pupil dilation response. For this analysis, pupil data were baseline-corrected to the average pupil size 2.5s prior to cue onset. This control analysis was added to increase sensitivity to detect potentially small reward-related pupil dilations that might otherwise be attenuated by overall task-evoked changes in pupil size.

To ensure that any observed differences were not driven by differences in the absolute baseline pupil diameter, we compared the average absolute pupil diameter during the final second of the baseline measurements using a paired t-test as a control analysis. Prior to analysis, normality of the data was assessed using the Shapiro-Wilk normality test.

In an exploratory, non-preregistered, control analysis, we examined whether the condition differences in pupil size observed during the tapping phase remained present during the break. Due to the shorter duration of the break (35s) compared to the tapping phase (40s), pupil data were binned into 5-second intervals (0–5s, 5–10s, 10–15s, 15–20s, 20–25s, 25–30s, 30–35s) instead of 10s intervals. A LMEM with the same model structure was fitted to the data using the previously described procedure.

#### Correlational analysis

To assess whether cue-evoked changes in pupil size were associated with cue-evoked changes in motor performance, we conducted correlational analyses on cue-evoked change scores for both conditions separately. For each variable, average values for the 5 seconds before and after cue onset were calculated. Change scores were derived by subtracting pre-cue from post-cue means for each variable. These cue-evoked change scores were subjected to a robust correlation analysis using the MATLAB-based Robust Correlation Toolbox^50^, which protects against bivariate outliers. For the reward condition, no bivariate outliers were detected and the assumption of multivariate normality was met, as assessed by the Henze-Zirkler test implemented in the Robust Correlation Toolbox. However, the data violated the assumption of homoscedasticity. For the neutral condition, a bivariate outlier was detected and the data violated both the assumption of normality and homoscedasticity. Consequently, we used Pearson correlation for the reward condition and Spearman correlation for the neutral condition and assessed significance using bootstrapped 95% confidence intervals, with effects considered significant if the interval excluded zero.

As an exploratory analysis, we compared the magnitude of the correlations of the cue-evoked changes between conditions (neutral vs. reward). To ensure comparability, the participant identified as an outlier in the neutral condition was removed from the reward dataset, after which the Spearman correlation for the reward condition was calculated. Correlations were statistically compared using the cocor package^51^ in R, which allows the comparison of two correlations obtained from the same cohort with non-overlapping variables.

## Results

### Reward increases motor performance in a fatigued state

We tested whether fatigability-related motor slowing was successfully induced by wrist tapping and whether it could be reduced by reward. Reward substantially modulated how motor performance evolved over time, as indicated by a significant effect of *Time × Condition* interaction effect (LMEM: *F*(3,144) = 7.676, *p* < 0.0001, 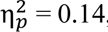, 95% CI for 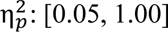; Figure 2), alongside a significant main effect of *Time* (*F*(3,144) = 7.044, *p* = 0.0002, 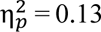, 95% CI for 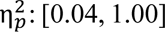: [0.04, 1.00]) and *Condition* (*F*(1,24) = 13.501, *p* = 0.001, 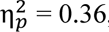, 95% CI for 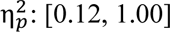). In the neutral condition, tapping frequency declined over time, with post hoc comparison revealing significantly lower tapping frequencies in the final two time bins (20-30s, 30-40s) compared to the initial 0-10s bin (*t* ≥ 3.925; *p* ≤ 0.0008), indicating that motor slowing was successfully induced. Within the reward condition, tapping frequency also decreased over time prior to the reward cue, even though this post-hoc comparison only reached trend-level significance after correcting for multiple comparisons (0-10s vs. 10-20s: *t*(144) = 2.631, *p* = 0.057). The reward cue presentation led to a significant increase in tapping speed: participants tapped faster in the first 10s after cue onset compared to the last 10s of the pre-cue phase (10-20s vs 20-30s: *p* < 0.0001). Importantly, participants tapped significantly faster in response to the reward cue (20-30s, 30-40s) compared to the neutral cue (|*t*| ≥ 4.379, *p* < 0.0001). No significant differences were observed between conditions prior to cue presentation (0-10s and 10-20s: *p* ≥ 0.607), indicating that both conditions were comparable before the reward cue was presented. The tapping speeds participants reached in response to the reward cue were not significantly different from their initial tapping speed (0-10s vs 20-30s: *p* = 1.00; 0-10s vs 30-40s: *p* = 1.00).

**Figure 2.**
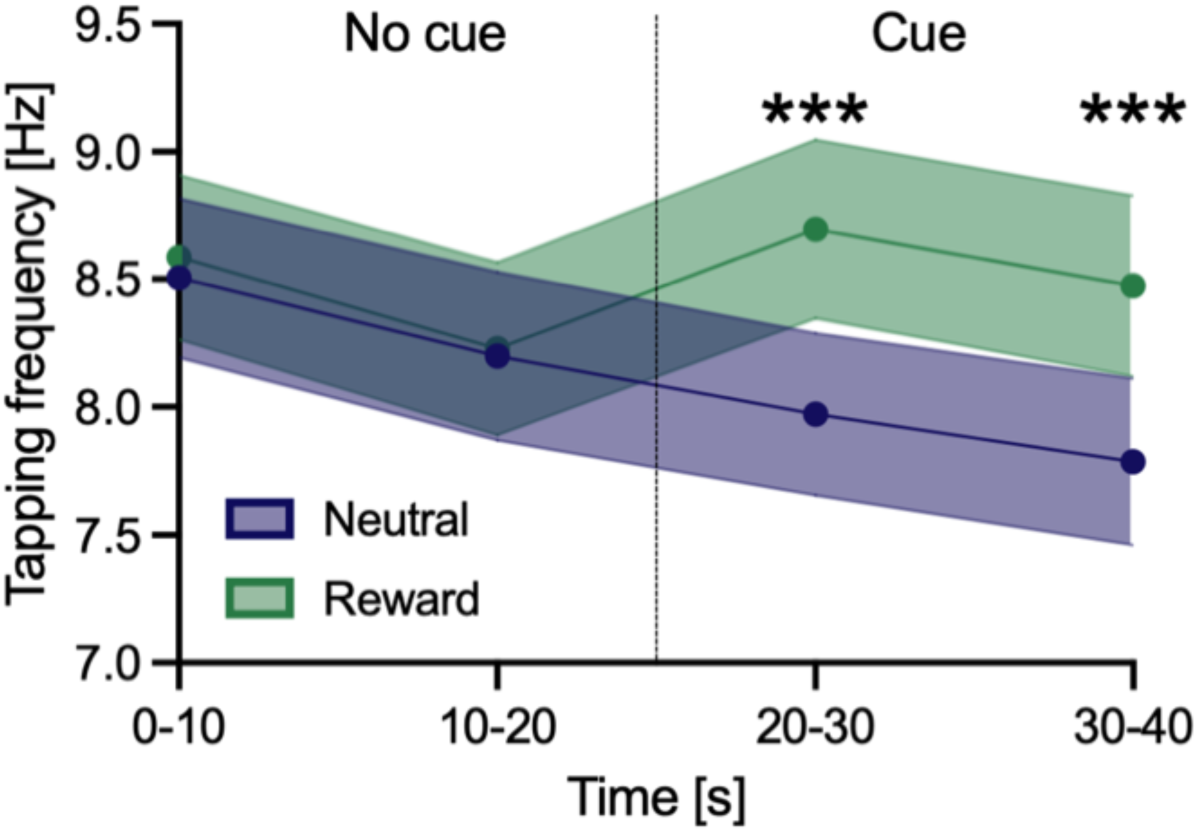
Behavioural results. Binned tapping frequency averaged across participants, for the neutral (blue) and reward (green) condition. Dashed vertical line indicates the cue onset after the first half of tapping (20s). Shaded areas represent the standard error of the mean (SEM). *** indicate significant differences between conditions with p < 0.001.

These findings indicate that participants were able to significantly increase their tapping speed when given the possibility to earn a reward, even though they were already entering a fatigued state. In response to reward, participants were able to reach tapping speeds comparable to their initial performance.

### Muscle activity increases over time

We investigated how muscle activity of the ECR and FCR, as well as their coactivation, changed over time during a motor slowing task and how reward modulated these patterns. Although the residuals of all three statistical models violated normality, (Kolmogorov-Smirnov test with *p* < 0.05), bootstrap validation supported the robustness of our parameter estimates, yielding confidence intervals highly consistent with model-based estimates (see *Methods Statistical analysis*). Detailed comparisons between model-based and bootstrap confidence intervals are provided in Supplementary Tables 1-3.

First, we examined the AUC of both the ECR and the FCR (Figure 3a). ECR activity increased over time, with a significant main effect of *Time* (*F*(3, 168) = 34.761, *p* < 0.0001, 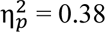, 95% CI for 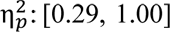). In contrast, no significant effect of reward on ECR activity was found (main effect of *Condition*: *p* = 0.264; *Time × Condition* interaction: *p* = 0.469). Given the absence of a *Time × Condition* interaction, post hoc comparisons addressing the main effect of *Time* were conducted on data pooled across conditions, revealing significantly higher muscle activity in all later time bins compared to the initial 0-10s bin (|*t*| ≥ = 3.355, *p* ≤ 0.006).

**Figure 3.**
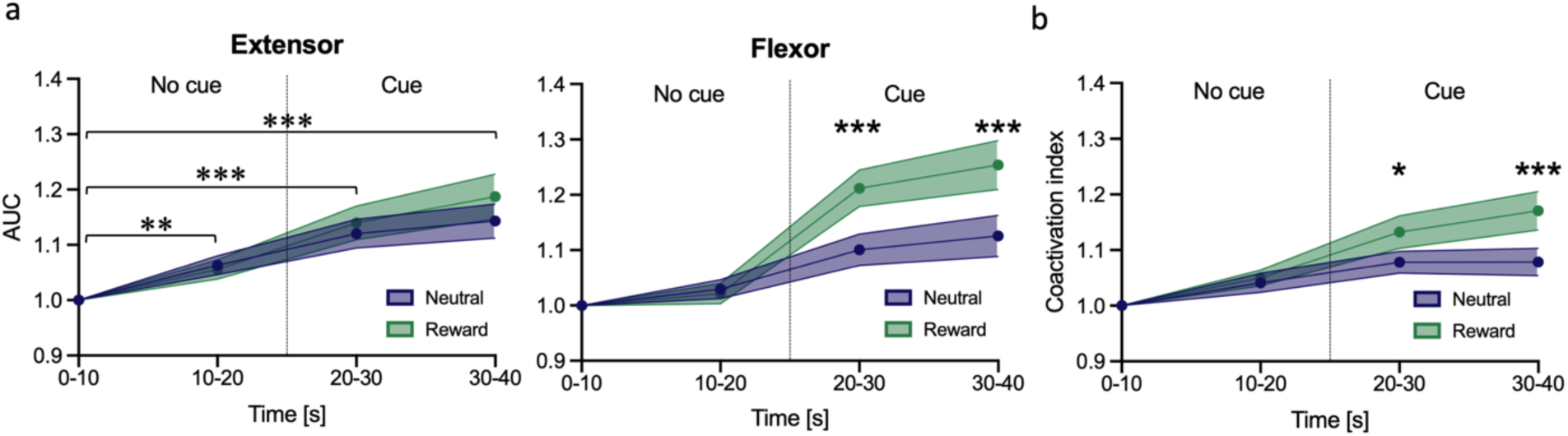
Area under the curve of the extensor carpi radialis longus (ECR) and flexor carpi radialis (FCR) muscles. Binned AUC normalized to the tapping speed and the AUC of bin 1 (0-10s) for the ECR (a, left) and FCR (a, right). (b) Coactivation index of the AUC of the ECR and FCR normalized to the AUC of bin 1(0-10s). Dashed vertical line indicates the cue onset after the first half of tapping (20s). Shaded areas represent the standard error of the mean (SEM). Brackets with asterisks (Panel a, left) indicate comparisons between each time bin (10-20s, 20-30s, 30-40s) and the first time bin (0-10s; pooled across condition). Asterisks indicate significant differences between time bins (panel a left) and conditions (panel a right; panel b). * p < .05, **p < .01, ***p < .001.

For the FCR, muscle activity also increased over time (main effects of *Time:* (*F*(3, 168) = 46.757, *p* < 0.0001, 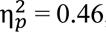, 95% CI for 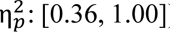), but unlike the ECR, it was also significantly modulated by the reward cue presentation (main effect *Condition*: *F*(1, 168) = 17.711, *p* < 0.0001, 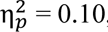, 95% CI for 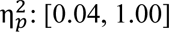: [0.04, 1.00]; *Time × Condition* interaction: *F*(3, 168) = 6.807, *p* = 0.0002, 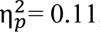 = 0.11, 95% CI for 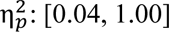: [0.04, 1.00]). In both conditions, the AUC of the FCR showed a similar pattern over time, with significant increases from 0-10s to 20-30s and to 30-40s (neutral: |*t*| ≥ 3.652, *p* ≤ 0.002; reward: |*t*| ≥ 7.691, *p* < 0.0001). However, reward significantly amplified this increase. Direct comparisons between condition revealed significantly higher AUC of the FCR in the reward compared to the neutral condition during the cue presentation phase (20-30s and 20-40s: |*t*| ≥ 4.039, *p* ≤ 0.0001), but not during the first half of tapping (0-10s and 10-20s: *p* ≥ 0.777). Within the reward condition, comparing the cue phase with the no-cue phase, muscle activity was significantly higher in the last two bins (20-30s, 30-40s) compared the last 10s of the no-cue phase (10-20s; |*t*| ≥ 6.907, *p* < 0.0001). These results demonstrate that while both extensor and flexor muscles showed increased activity over time during the motor slowing task, only the flexor muscle exhibited significant reward-related modulation.

To better understand how the coactivation pattern of this antagonistic muscle pair changes over time and in response to reward, we looked at the coactivation index of the ECR and FCR muscles (Figure 3b). This coactivation index changed over time (main effect of *Time*: *F*(3, 144) = 26.566, *p* < 0.0001, 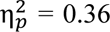, 95% CI for 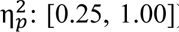 and was modulated by reward, as indicated by a significant main effect of *Condition* (*F*(1, 24) = 7.922, *p* = 0.010, 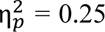, 95% CI for 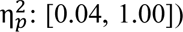) and a significant interaction effect of *Time x Condition* (*F*(3, 144) = 3.735, *p* = 0.013, 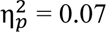, 95% CI for 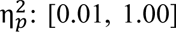). Coactivation indices were significantly higher in later time bins (20-30s, 30-40s) compared to 0-10s in both the neutral and reward condition (neutral: |*t*| ≥ 3.543, *p* ≤ 0.032; reward: |*t*| ≥ 5.995, *p* < 0.0001). The *Time x Condition* interaction was driven by a stronger increase in the coactivation index after reward-cue presentation compared to the neutral cue (20-30s, 30-40s: |*t*| ≥ 2.298, *p* ≤ 0.023), while no significant difference was detected prior to cue presentation (10-20s: *p* = 0.707). Overall, these results indicate that the ECR and the FCR were increasingly coactivated over the time course of the trial with this coactivation pattern being further enhanced in response to the reward cue.

### Pupil size increases in response to the reward cue

To investigate how pupil size changed during motor slowing and in response to reward, we conducted a statistical analysis of the binned pupil data. Pupil size was modulated by time and rewards as indicated by a significant main effect of *Time* (LMEM: *F*(3,144) = 13.577, *p* < 0.0001, 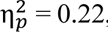 = 0.22, 95% CI for 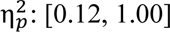: [0.12, 1.00]), *Condition* (*F*(1,24) = 10.661, *p* = 0.003, 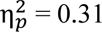 = 0.31, 95% CI for 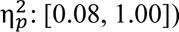: [0.08, 1.00]), and their interaction (*Time × Condition:* (*F*(3,144) = 14.077, *p* < 0.0001, 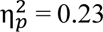 = 0.23, 95% CI for 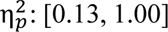: [0.13, 1.00]). Post hoc analysis revealed that in the neutral condition pupil size significantly decreased during the last three time bins compared to the first bin (*t* ≥ 4.364, *p* ≤ 0.0001; Supplementary Figure 2b). In the reward condition, this constriction pattern was disrupted by a cue-evoked pupil dilation. Comparable to the neutral condition, pupil size initially decreased from the 0-10s to 10-20s bin (*t*(144) = 4.47, *p* = 0.0001). However, following reward cue presentation at 20s, pupil size increased significantly compared to the preceding time bin (10-20s vs 20-30s and 10-20s vs 30-40s: |*t*| ≥ 4.570, *p* ≤ 0.0001), returning to similar pupil sizes observed during the initial pupil dilation (0-10s vs 20-30s: *p* = 0.250; 0-10s vs 30-40s: *p* = 1.00). Post hoc tests comparing pupil size between the reward and neutral condition revealed significantly larger pupil sizes in the reward condition during the 20-30s and the 30-40s time bins (|*t*| ≥ 4.070, *p* < 0.0001).

In addition to this pre-registered analysis, we added tapping speed as a covariate to our LMEM to explore whether pupil size explains variance beyond that attributable to faster tapping per se. As expected, tapping speed itself had a significant effect on pupil size (*Tapping Frequency*: *F*(3,119.955) = 13.452, *p* = 0.0003, 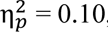 95% CI for 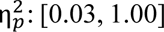). However, the previously described effects remained significant (*Time*: *F*(3,145.060) = 10.917, *p* < 0.0001, 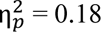, 95% CI for 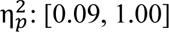; *Condition*: *F*(3,25.206) = 7.368, *p* = 0.012, 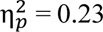, 95% CI for 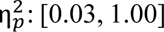; *Time x Condition*: *F*(3,145.348) = 9.960, *p* < 0.0001, 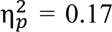, 95% CI for 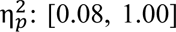). This indicates that the observed changes in pupil size are not solely attributable to the fact that participants tapped faster in the reward condition or at the beginning of the trial.

To more precisely identify when pupil size differed between the two conditions, we statistically compared the continuous, baseline-corrected pupil size time-series (Figure 4a) using SPM1D. As expected, pupil trajectories did not differ significantly for the first half of the tapping phase, prior to cue onset. However, following cue onset (i.e., after first 20s of tapping phase), pupil size significantly differed between conditions, with a larger pupil dilation observed in the reward condition (SPM{z*} = 3.535, 5 supra threshold clusters with the largest cluster p = 0.017, smallest cluster p < 0.001; black line in Figure 4a indicating time points with statistically significant condition effect).

**Figure 4.**
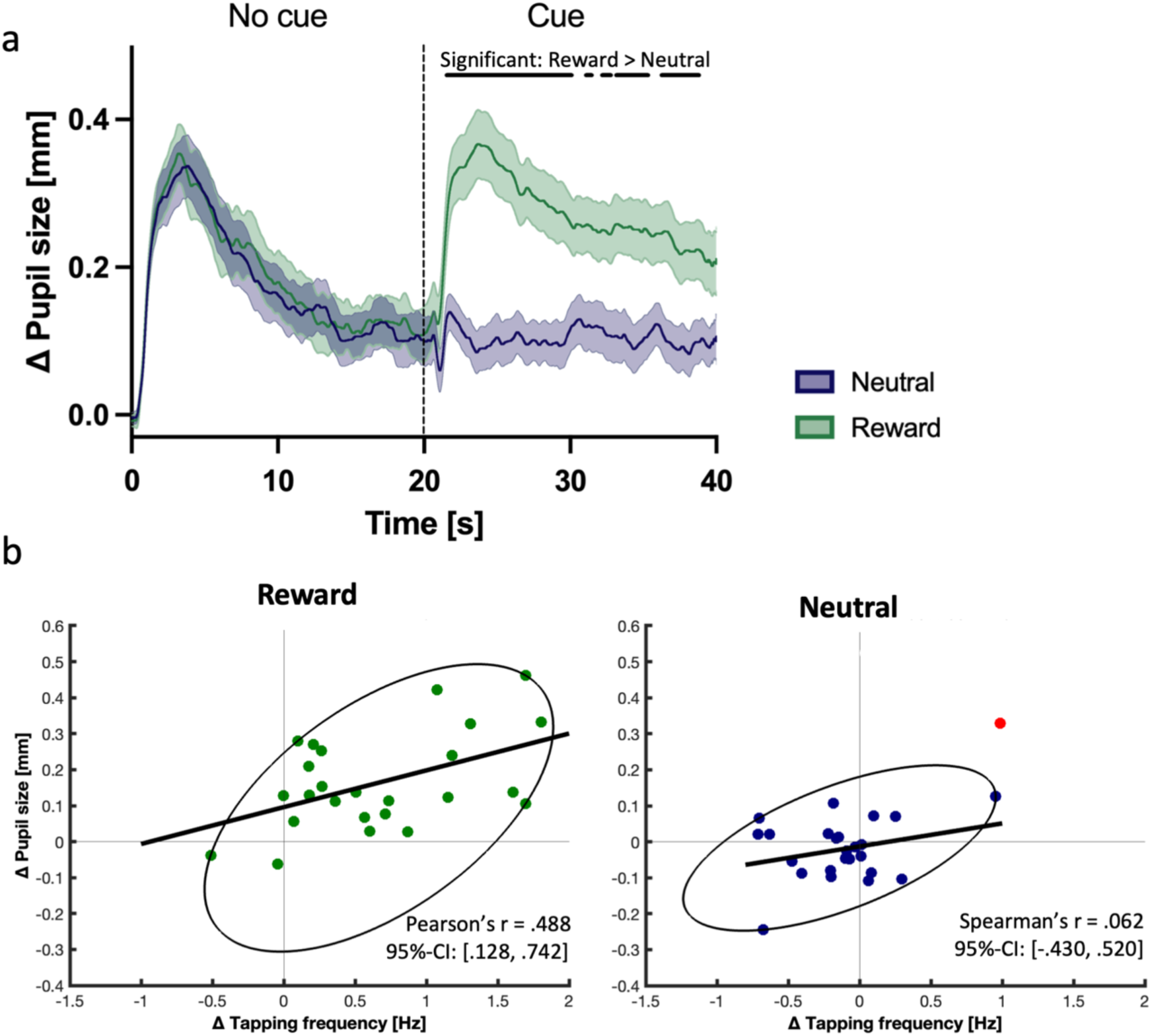
Pupil results and correlations of cue-evoked changes. (a) Baseline-corrected pupil diameter, averaged across participants, for the neutral (blue) and reward (green) condition. Dashed vertical lines indicate the cue onset after the first half of tapping (20s). Shaded areas represent the standard error of the mean (SEM). Black lines/dots denote time clusters showing significant differences (p ≤ 0.05) between conditions (reward vs. neutral). Correlations between cue-evoked changes in tapping frequency and pupil size, calculated as the difference between the mean signal in the 5s post-cue and 5s pre-cue periods, shown separately for the reward (b) and neutral (c) condition. Each dot represents one participant, red dots indicate statistical outliers excluded from the correlation analysis, and ellipses denote the data range included in the correlation.

### Reward-cue evoked changes in tapping speed correlate with changes in pupil size

To examine whether reward cue-evoked changes in pupil size and tapping speed were positively correlated, we computed a robust correlation with the reward cue-evoked change scores. Reward cue-evoked changes in tapping frequency and pupil size positively correlated (Pearson’s *r* = 0.488, 95% CI: [0.128, 0.742]; Figure 4b). We additionally included the corresponding correlation for the neutral condition, in which participants were also instructed to tap as fast as possible. The correlation for the neutral condition was close to zero (Spearman’s ρ = 0.062, 95% CI: [-0.430, 0.520]), consistent with the absence of a cue-evoked behavioural change in this condition. To directly test whether the association between tapping speed and pupil size was stronger in the reward condition than in the neutral condition, we conducted an exploratory between-condition comparison. We computed Spearman’s ρ for both conditions (after removing a neutral-condition outlier from both datasets). This analysis revealed no significant difference in correlation strength (reward: Spearman’s ρ = 0.326, 95% CI: [-0.176, 0.693]; ρ Spearman’s ρ = 0.264, 95% CI: [-0.762, 0.280]). The absence of a significant difference may reflect limited statistical power, as the study was not specifically designed to test between-condition differences in correlation strength.

### Control analyses

To confirm that the baseline correction applied to the pupil data did not introduce systematic differences between conditions, we compared the average absolute pupil diameters of the last second of the baseline period (Supplementary Figure 2a). The data was normally distributed (Shapiro-Wilk test: *p* = 0.752), and a paired-samples t-test revealed no significant differences in baseline pupil diameters between conditions (*t*(24) = 1.57, *p* = 0.130). Thus, we found no statistical evidence suggesting that the average baseline pupil size differed across conditions or that it did drive the above-described effects.

We also conducted a preregistered analysis comparing cue-evoked pupil responses using pupil time series corrected to the average pupil size 2.5 s prior to cue onset. We preregistered this analysis, as we anticipated more subtle changes in pupil size. This analysis supports our main analysis but did not yield new insights, as the main time series analysis already revealed a robust and visually evident reward-related pupil dilation. Full results are available in the Supplementary Figure 2c.

Finally, we performed an exploratory analysis that examined pupil dynamics during the break period following the tapping task to explore whether differences observed between conditions remain during the break (Supplementary Figure 2e). Across conditions, we observed a progressive decrease in pupil size over time (LMEM main effect of *Time*, *F*(6,288) = 32.861, *p* < 0.0001, 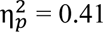 = 0.41, 95% CI for 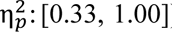, with significantly smaller pupil sizes during all time bins of the break compared to the initial 5s bin of the break (neutral: all *p* ≤ 0.009; reward: all *p* ≤ 0.0001). Neither *Condition* nor the *Time x Condition* interaction reached significance (main effect of *Condition*: *F*(1,24) = 2.526, *p* = 0.146, 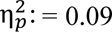= 0.09, 95% CI for 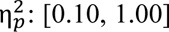: [0.10, 1.00]; interaction effect *Time x Condition*: *F*(6,288) = 0.933, *p* = 0.471, 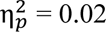 = 0.02, 95% CI for 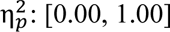: [0.00, 1.00]). As model residuals violated normality (Kolmogorov-Smirnov test, *p* < 0.05), we conducted bootstrap validation, which broadly supported our estimates but yielded diverging confidence intervals for the *Time × Condition* interaction (see Supplementary Table 4). Given this discrepancy and the exploratory nature of the analysis, these findings should be interpreted with caution.

## Discussion

In this study, we investigated whether the possibility to earn a reward can reduce fatigability-related motor slowing, and how muscle activity and pupil dilation change in response to it. Our findings demonstrate that the possibility to earn a reward led to a voluntary increase in tapping speed, even though participants were already in a fatigued state. However, this increase in motor performance was accompanied by higher muscle (co-)activation indicating that moving became more “costly” compared to the non-fatigued state. For pupil size, we observed an initial dilation at movement onset followed by a gradual decrease over time, and an increase in response to the reward cue. Reward-cue evoked changes in pupil size positively correlated with the reward-cue evoked changes in tapping speed, although the strength of this relationship was not significantly greater than that observed in the neutral condition. Pupil size changes were stronger than tapping speed alone could account for, suggesting that the observed effects cannot be fully explained by increases in tapping speed.

Our results provide evidence that reward attenuates motor slowing. Specifically, the presentation of the reward cue led to a significant increase in tapping speed compared to the neutral cue, even when participants were already in a fatigued state. Previous studies have established that reward enhances motor performance in a non-fatigued state^10,11,15–17,19^. The present findings expand on this literature by demonstrating that the performance-enhancing effects of reward persist even when being in a fatigued state, helping to counteract the decline in tapping speed that characterises motor slowing. This highlights that motor fatigability can be modulated by motivational state.

Given that the performance-enhancing effect of reward persists in a fatigued state, the question arises how reward mitigates motor fatigability. Previous work in a non-fatigued state showed that reward can enhance motor performance either by optimizing motor control^10,11,16,17^ or by the invigoration of movements^15,17,19^. In the context of motor fatigability, muscle activity patterns can offer indirect insight into the efficiency of movement execution. Previous work has demonstrated that coactivation of an antagonistic muscle pair increases during a motor slowing task, which might result from a breakdown in surround inhibition^3^ or general disinhibition of the sensorimotor cortex^5^. Thus, motor slowing is associated with an already inefficient motor control state, in which higher coactivation reflects more energetically costly movement execution. Consistent with this framework, we observed higher AUC of the ECR and FCR muscles and their coactivation per tap over the course of the motor slowing task. Crucially, reward did not reverse this pattern but amplified it: AUC of the FCR and coactivation index of the ECR and FCR muscle were significantly higher during reward cue compared to the neutral cue presentation. Thus, rather than becoming more efficient with reward, muscle activity became energetically even more demanding. The voluntary, reward-induced increase in tapping speed was therefore achieved at the expense of higher energetic costs per movement. One possible interpretation is that participants may have exerted greater force on the flexion side in response to the reward cue, potentially accepting less efficient and more effortful motor execution in exchange for performance gains. This pattern is in accordance with Shadmehr et al.^52^, who showed that reward can justify greater energetic cost. Hence, our EMG results indicate that reward mitigates motor slowing by invigorating motor output at increased energetic cost, rather than optimizing motor control.

To better contextualize the observed muscle activity patterns, it is important to consider potential contributions of both central and peripheral mechanisms to motor slowing. Converging evidence suggests that motor slowing is strongly driven by supraspinal mechanisms, as peripheral or spinal mechanisms such as maximal voluntary contraction^4,9^, responses to peripherals stimulation^8,9,53^, voluntary activation^9^, spinal excitability^53^ and inhibition^8^ remain relatively stable, whereas corticomotor excitability^9^ and intracortical inhibitory processes within sensorimotor cortex show pronounced changes^3,7,53^. However, these studies were conducted using repetitive finger tapping paradigms engaging small finger muscles. In the present study, we also used a repetitive tapping paradigm, but we used wrist tapping instead of finger tapping. Since larger muscles contribute to wrist tapping, it is possible that there is a stronger peripheral contribution compared to finger tapping, as the relative contribution of these mechanisms to motor fatigability is task dependent^6,8^. Force-based paradigms engage peripheral mechanisms more strongly, but even when using force paradigms, motor fatigability has been shown to be at least partly associated with suboptimal central drive^54–56^. Within this framework, during fatigability-inducing wrist tapping, the central motor drive may become suboptimal and reward may enable participants to tap into a performance reserve, allowing them to transiently mobilize more resources leading to an increase in central drive onto an already inefficient motor output. As we did not directly assess neural activity, these interpretations remain speculative and will require future studies using neuroimaging or brain stimulation techniques.

We examined how motor slowing and reward affect pupil size. For neutral tapping trials we observed a pronounced pupil dilation when tapping was initiated, followed by a gradual decrease in pupil size over time. These time series characteristics are typical for pupil responses and can be modelled with an impulse response function^57–59^. Responding to the reward cue caused similar pupil dynamics but both the initial peak and the final plateau of pupil size were significantly larger than in the neutral condition. These findings replicate and complement previous results linking larger pupil dilation to higher physical effort for discrete movements, which typically cause a strong phasic dilation response^23–26^, and for sustained movements, which additionally cause prolonged pupil dilation^22^. Similarly, pupil size appears to also reflect anticipated effort when tested with an effort-based decision making task, especially when participants accepted the physical effort compared to its avoidance^60^. In these paradigms, higher physical effort was operationalized as higher force, number of movements, amplitude or cycling intensity requirements. These results indicate a general sensitivity of the pupil to physical demand. In our data, tapping speed significantly affected pupil size, consistent with these prior observations. However, controlling for tapping speed as a covariate, the observed effects of *Time*, *Condition*, and their interaction on pupil size became smaller but remained significant. Thus, changes in pupil size in response to the reward cue cannot be explained by tapping speed per se, which, according to our instructions, was the primary variable that needed to be controlled by the participants. Given that fatigability affects the efficiency of the motor system such that higher EMG activity is required per tap, the sustained pupil dilation in response to the reward cue likely reflects that reward motivated higher effort investment for generating higher tapping speed at increased energetic cost. An alternative interpretation is that pupil dilation exceeding that predicted by tapping speed reflects reward processing itself. Indeed, previous research has shown larger pupil sizes accompanied by shorter reaction times during high-reward compared to low- or no-reward trials^28,29^. However, the cooccurrence of pupil dilation with faster motor responses in these studies already suggests that the pupil is not tracking reward processing in isolation, but rather a state that benefits task performance more broadly. Consistent with this, pupil dilations during reward anticipation have been associated with increased BOLD activity in the salience network, which is thought to mediate an increase in arousal to improve task performance^29^. Within the context of our paradigm, we therefore argue that reward and effort investment are closely linked to arousal mechanisms mediated by cholinergic and, particularly, noradrenergic activity^61–64^. The locus coeruleus noradrenaline system (LC-NA) is a key neuromodulatory brain system involved in regulating the brain’s arousal level^63,65^. Importantly, LC-NA activity causally influences pupil size^66–68^. The LC-NA system has been implicated in mobilizing resources required for goal-directed behaviour. For example, in non-human primates, increased physical effort in response to reward has been associated with heightened LC activity, and pupil dilations scale with firing of noradrenergic neurons in the LC^24,69^. Pharmacological manipulations further support a causal role: reducing noradrenaline availability with clonidine diminishes force production^70,71^. Notably, the LC–NA system does not operate in isolation but interacts with dopaminergic circuits involved in reward processing^24^, suggesting that motivational cues may engage coordinated neuromodulatory mechanisms to support sustained effort. Since pupil size under constant lighting conditions can be used to approximate the activity of the LC-NA system^72,73^, our findings provide converging, albeit indirect evidence that LC-NA activity may support effort mobilization in response to reward cues, even as motor fatigability progressively reduces the efficiency of movement execution and increases its energetic cost.

While pupil size is widely used to approximate the activity of the LC-NA system^72,73^, it is important to acknowledge that other neuromodulatory systems, such as the cholinergic system, may also be potential contributors^74^. Future studies employing pharmacological manipulations or neuromelanin-sensitive MRI sequences in combination with fMRI could provide more direct insights into the function of the LC-NA system in motor performance under reward and motor fatigability in humans.

In summary, reward attenuates motor fatigability as measured via tapping speed but we found no evidence that this has been achieved by restoring efficient motor control. Instead, our data suggests that it enables individuals to mobilize additional resources to counteract fatigability most likely by accepting higher movement costs as indicated by increased muscle activation per tap. This is associated with greater effort investment, as indicated by sustained pupil dilation, which cannot be solely explained by tapping speed. These findings suggest that motivational interventions such as reward could be leveraged in populations with impaired motor fatigability, including neurological rehabilitation settings.

## Data availability statement

The data supporting the findings of this study will be openly available in the ETH Zurich Research Collection.

## Acknowledgments

The authors would like to thank all participants for their involvement in the study. We are especially grateful to Nathalie Buchmann for her assistance with data collection, Dr. Daniel Woolley for his technical support, and Dr. Snow (Xue) Zhang for her valuable contributions throughout the project. We also thank Paige Howell for proof-reading, Eduardo Villar for the drawing of the experimental setup, and the Seminar for Statistics at ETH Zurich for statistical consulting. This work was supported by the Swiss National Science Foundation (SNSF) grant 32003B_207719. S.N.M is supported by the SNSF starting grant TMSGI1_226129. N.W. is supported by the National Research Foundation, Prime Minister’s Office, Singapore under its Campus for Research Excellence and Technological Enterprise (CREATE) programme.

## Author contributions

J.I., C.H., M.B., S.N.M., R.R., and N.W. were involved in conceptualization and design of the study. J.I. programmed the task and the analysis scripts and acquired the data. J.I., C.H., M.B., R.R., and N.W. planned the analysis. J.I. analysed the data. J.I., C.H., M.B., S.N.M., R.R., and N.W. interpreted the data. J.I. drafted the manuscript and all authors substantively revised it.

## Conflict of interest

All authors declare no competing interests.

## Notes

### Competing Interest Statement

The authors have declared no competing interest.

## References

1. Kluger, B. M., Krupp, L. B. & Enoka, R. M. Fatigue and fatigability in neurologic illnesses. Neurology 80, 409–416 (2013).

2. Manjaly, Z.-M. et al. Pathophysiological and cognitive mechanisms of fatigue in multiple sclerosis. J. Neurol. Neurosurg. Psychiatry 90, 642–651 (2019).

3. Bächinger, M. et al. Human motor fatigability as evoked by repetitive movements results from a gradual breakdown of surround inhibition. eLife 8, e46750 (2019).

4. Rodrigues, J. P., Mastaglia, F. L. & Thickbroom, G. W. Rapid slowing of maximal finger movement rate: fatigue of central motor control? Exp. Brain Res. 196, 557–563 (2009).

5. Heimhofer, C. et al. Dynamic causal modelling highlights the importance of decreased self-inhibition of the sensorimotor cortex in motor fatigability. Brain Struct. Funct. https://doi.org/10.1007/s00429-024-02840-1 (2024) doi:10.1007/s00429-024-02840-1.

6. Arias, P. et al. Central fatigue induced by short-lasting finger tapping and isometric tasks: A study of silent periods evoked at spinal and supraspinal levels. Neuroscience 305, 316–327 (2015).

7. Madinabeitia-Mancebo, E., Madrid, A., Oliviero, A., Cudeiro, J. & Arias, P. Peripheral-central interplay for fatiguing unresisted repetitive movements: a study using muscle ischaemia and M1 neuromodulation. Sci. Rep. 11, 2075 (2021).

8. Madrid, A., Valls-Solé, J., Oliviero, A., Cudeiro, J. & Arias, P. Differential responses of spinal motoneurons to fatigue induced by short-lasting repetitive and isometric tasks. Neuroscience 339, 655–666 (2016).

9. Madrid, A., Madinabeitia-Mancebo, E., Cudeiro, J. & Arias, P. Effects of a Finger Tapping Fatiguing Task on M1-Intracortical Inhibition and Central Drive to the Muscle. Sci. Rep. 8, 9326 (2018).

10. Codol, O., Holland, P. J., Manohar, S. G. & Galea, J. M. Reward-Based Improvements in Motor Control Are Driven by Multiple Error-Reducing Mechanisms. J. Neurosci. 40, 3604–3620 (2020).

11. Manohar, S. G., Muhammed, K., Fallon, S. J. & Husain, M. Motivation dynamically increases noise resistance by internal feedback during movement. Neuropsychologia 123, 19–29 (2019).

12. Opris, I., Lebedev, M. & Nelson, R. Motor Planning under Unpredictable Reward: Modulations of Movement Vigor and Primate Striatum Activity. Front. Neurosci. 5, (2011).

13. Pessiglione, M. et al. How the brain translates money into force: a neuroimaging study of subliminal motivation. Science 316, 904–906 (2007).

14. Pessiglione, M. et al. Subliminal Instrumental Conditioning Demonstrated in the Human Brain. Neuron 59, 561–567 (2008).

15. Summerside, E. M., Shadmehr, R. & Ahmed, A. A. Vigor of reaching movements: reward discounts the cost of effort. J. Neurophysiol. 119, 2347–2357 (2018).

16. Takikawa, Y., Kawagoe, R., Itoh, H., Nakahara, H. & Hikosaka, O. Modulation of saccadic eye movements by predicted reward outcome. Exp. Brain Res. 142, 284–291 (2002).

17. Manohar, S. G. et al. Reward Pays the Cost of Noise Reduction in Motor and Cognitive Control. Curr. Biol. 25, 1707–1716 (2015).

18. Marbaker, R. M., Schmad, R. C., Al-Ghamdi, R. A., Sukumar, S. & Ahmed, A. A. Reward invigorates isometric gripping actions. J. Neurophysiol. 133, 1282–1294 (2025).

19. Shadmehr, R., Reppert, T. R., Summerside, E. M., Yoon, T. & Ahmed, A. A. Movement Vigor as a Reflection of Subjective Economic Utility. Trends Neurosci. 42, 323–336 (2019).

20. Dudman, J. T. & Krakauer, J. W. The basal ganglia: from motor commands to the control of vigor. Curr. Opin. Neurobiol. 37, 158–166 (2016).

21. van der Wel, P. & van Steenbergen, H. Pupil dilation as an index of effort in cognitive control tasks: A review. Psychon. Bull. Rev. 25, 2005–2015 (2018).

22. Hayashi, N., Someya, N. & Fukuba, Y. Effect of Intensity of Dynamic Exercise on Pupil Diameter in Humans. J. Physiol. Anthropol. 29, 119–122 (2010).

23. Richer, F. & Beatty, J. Pupillary Dilations in Movement Preparation and Execution. Psychophysiology 22, 204–207 (1985).

24. Varazzani, C., San-Galli, A., Gilardeau, S. & Bouret, S. Noradrenaline and dopamine neurons in the reward/effort trade-off: a direct electrophysiological comparison in behaving monkeys. J. Neurosci. Off. J. Soc. Neurosci. 35, 7866–7877 (2015).

25. Voudouris, D., Schuetz, I., Schinke, T. & Fiehler, K. Pupil dilation scales with movement distance of real but not of imagined reaching movements. J. Neurophysiol. 130, 104–116 (2023).

26. Zénon, A., Sidibé, M. & Olivier, E. Pupil size variations correlate with physical effort perception. Front. Behav. Neurosci. 8, (2014).

27. Nieuwenhuis, S., De Geus, E. J. & Aston-Jones, G. The anatomical and functional relationship between the P3 and autonomic components of the orienting response. Psychophysiology 48, 162–175 (2011).

28. Cole, L., Lightman, S., Clark, R. & Gilchrist, I. D. Tonic and phasic effects of reward on the pupil: implications for locus coeruleus function. Proc. R. Soc. B Biol. Sci. 289, 20221545 (2022).

29. Schneider, M., Leuchs, L., Czisch, M., Sämann, P. G. & Spoormaker, V. I. Disentangling reward anticipation with simultaneous pupillometry / fMRI. NeuroImage 178, 11–22 (2018).

30. Hartmann, E. V., Reichert, C. F. & Spitschan, M. Effects of caffeine intake on pupillary parameters in humans: a systematic review and meta-analysis. Behav. Brain Funct. 20, 19 (2024).

31. Niehorster, D. C., Andersson, R. & Nyström, M. Titta: A toolbox for creating PsychToolbox and Psychopy experiments with Tobii eye trackers. Behav. Res. Methods 52, 1970–1979 (2020).

32. Brainard, D. H. The Psychophysics Toolbox. Spat. Vis. 10, 433–436 (1997).

33. Kleiner, M. et al. What’s new in Psychtoolbox-3? Perception 36, 1–16 (2007).

34. Pelli, D. G. The VideoToolbox software for visual psychophysics: transforming numbers into movies. Spat. Vis. 10, 437–442 (1997).

35. De Luca, C. J., Donald Gilmore, L., Kuznetsov, M. & Roy, S. H. Filtering the surface EMG signal: Movement artifact and baseline noise contamination. J. Biomech. 43, 1573–1579 (2010).

36. Frost, G., Dowling, J., Dyson, K. & Bar-Or, O. Cocontraction in three age groups of children during treadmill locomotion. J. Electromyogr. Kinesiol. 7, 179–186 (1997).

37. Kret, M. E. & Sjak-Shie, E. E. Preprocessing pupil size data: Guidelines and code. Behav. Res. Methods 51, 1336–1342 (2019).

38. Mathôt, S., Fabius, J., Van Heusden, E. & Van der Stigchel, S. Safe and sensible preprocessing and baseline correction of pupil-size data. Behav. Res. Methods 50, 94–106 (2018).

39. Fink, L. et al. From pre-processing to advanced dynamic modeling of pupil data. Behav. Res. Methods 56, 1376–1412 (2024).

40. Bates, D., Mächler, M., Bolker, B. & Walker, S. Fitting Linear Mixed-Effects Models Using lme4. J. Stat. Softw. 67, 1–48 (2015).

41. Barr, D. J., Levy, R., Scheepers, C. & Tily, H. J. Random effects structure for confirmatory hypothesis testing: Keep it maximal. J. Mem. Lang. 68, (2013).

42. Hartig, F. DHARMa: Residual Diagnostics for Hierarchical (Multi-Level / Mixed) Regression Models. (2024).

43. Loy, A., Steele, S. & Korobova, J. lmeresampler: Bootstrap Methods for Nested Linear Mixed-Effects Models. (2025).

44. Kuznetsova, A., Brockhoff, P. B. & Christensen, R. H. B. lmerTest: Tests in Linear Mixed Effects Models. Journal of Statistical Software 82, 1--26 (2017).

45. Ben-Shachar, M., Lüdecke, D. & Makowski, D. effectsize: Estimation of Effect Size Indices and Standardized Parameters}. Journal of Open Source Software 5, 2815 (2020).

46. Lenth, R. V. emmeans: Estimated Marginal Means, aka Least-Squares Means. (2024).

47. Pataky, T. C., Robinson, M. A. & Vanrenterghem, J. Vector field statistical analysis of kinematic and force trajectories. J. Biomech. 46, 2394–2401 (2013).

48. Pataky, T. C., Vanrenterghem, J. & Robinson, M. A. Zero- vs. one-dimensional, parametric vs. non-parametric, and confidence interval vs. hypothesis testing procedures in one-dimensional biomechanical trajectory analysis. J. Biomech. 48, 1277–1285 (2015).

49. Meissner, S. N. et al. Self-regulating arousal via pupil-based biofeedback. *Nat*. Hum. Behav. 1–20 (2023) doi:10.1038/s41562-023-01729-z.

50. Pernet, C. R., Wilcox, R. R. & Rousselet, G. A. Robust Correlation Analyses: False Positive and Power Validation Using a New Open Source Matlab Toolbox. Front. Psychol. 3, (2013).

51. Diedenhofen, B. Comparing Correlations. (2022).

52. Shadmehr, R., Huang, H. J. & Ahmed, A. A. A Representation of Effort in Decision-Making and Motor Control. Curr. Biol. 26, 1929–1934 (2016).

53. Madinabeitia-Mancebo, E., Madrid, A., Jácome, A., Cudeiro, J. & Arias, P. Temporal dynamics of muscle, spinal and cortical excitability and their association with kinematics during three minutes of maximal-rate finger tapping. Sci. Rep. 10, 3166 (2020).

54. Gandevia, S. C., Allen, G. M., Butler, J. E. & Taylor, J. L. Supraspinal factors in human muscle fatigue: evidence for suboptimal output from the motor cortex. J. Physiol. 490, 529–536 (1996).

55. Smith, J. L., Martin, P. G., Gandevia, S. C. & Taylor, J. L. Sustained contraction at very low forces produces prominent supraspinal fatigue in human elbow flexor muscles. J. Appl. Physiol. Bethesda Md 1985 103, 560–568 (2007).

56. Søgaard, K., Gandevia, S. C., Todd, G., Petersen, N. T. & Taylor, J. L. The effect of sustained low-intensity contractions on supraspinal fatigue in human elbow flexor muscles. J. Physiol. 573, 511–523 (2006).

57. Abivardi, A. et al. Acceleration of inferred neural responses to oddball targets in an individual with bilateral amygdala lesion compared to healthy controls. Sci. Rep. 13, 14550 (2023).

58. Korn, C. W., Staib, M., Tzovara, A., Castegnetti, G. & Bach, D. R. A pupil size response model to assess fear learning. Psychophysiology 54, 330–343 (2017).

59. Korn, C. W. & Bach, D. R. A solid frame for the window on cognition: Modelling event-related pupil responses. J. Vis. 16, 28 (2016).

60. Kurniawan, I. T., Grueschow, M. & Ruff, C. C. Anticipatory Energization Revealed by Pupil and Brain Activity Guides Human Effort-Based Decision Making. J. Neurosci. 41, 6328–6342 (2021).

61. Jones, B. E. Activity, modulation and role of basal forebrain cholinergic neurons innervating the cerebral cortex. in Progress in Brain Research vol. 145 157–169 (Elsevier, 2004).

62. Jones, B. E. Arousal systems. Front. Biosci. 8, 438–451 (2003).

63. Aston-Jones, G. & Cohen, J. D. An integrative theory of locus coeruleus-norepinephrine function: adaptive gain and optimal performance. Annu. Rev. Neurosci. 28, 403–450 (2005).

64. Hasselmo, M. E. & Sarter, M. Modes and Models of Forebrain Cholinergic Neuromodulation of Cognition. Neuropsychopharmacology 36, 52–73 (2011).

65. Robertson, S. D., Plummer, N. W., de Marchena, J. & Jensen, P. Developmental origins of central norepinephrine neuron diversity. Nat. Neurosci. 16, 1016–1023 (2013).

66. Hayat, H. et al. Locus coeruleus norepinephrine activity mediates sensory-evoked awakenings from sleep. Sci. Adv. 6, eaaz4232 (2020).

67. Megemont, M., McBurney-Lin, J. & Yang, H. Pupil diameter is not an accurate real-time readout of locus coeruleus activity. eLife 11, e70510 (2022).

68. Zerbi, V. et al. Rapid Reconfiguration of the Functional Connectome after Chemogenetic Locus Coeruleus Activation. Neuron 103, 702–718.e5 (2019).

69. Bouret, S. & Richmond, B. J. Sensitivity of Locus Ceruleus Neurons to Reward Value for Goal-Directed Actions. J. Neurosci. 35, 4005–4014 (2015).

70. Borderies, N., Bornert, P., Gilardeau, S. & Bouret, S. Pharmacological evidence for the implication of noradrenaline in effort. PLOS Biol. 18, e3000793 (2020).

71. Jahn, C. I. et al. Dual contributions of noradrenaline to behavioural flexibility and motivation. Psychopharmacology (Berl.) 235, 2687–2702 (2018).

72. Murphy, P. R., Robertson, I. H., Balsters, J. H. & O’connell, R. G. Pupillometry and P3 index the locus coeruleus–noradrenergic arousal function in humans. Psychophysiology 48, 1532–1543 (2011).

73. Murphy, P. R., O’Connell, R. G., O’Sullivan, M., Robertson, I. H. & Balsters, J. H. Pupil diameter covaries with BOLD activity in human locus coeruleus. Hum. Brain Mapp. 35, 4140–4154 (2014).

74. Reimer, J. et al. Pupil fluctuations track rapid changes in adrenergic and cholinergic activity in cortex. Nat. Commun. 7, 13289 (2016).

