## Supplementary Material for "Reward Reduces Motor Fatigability by Increasing Movement Vigour"

#### Supplementary Figures

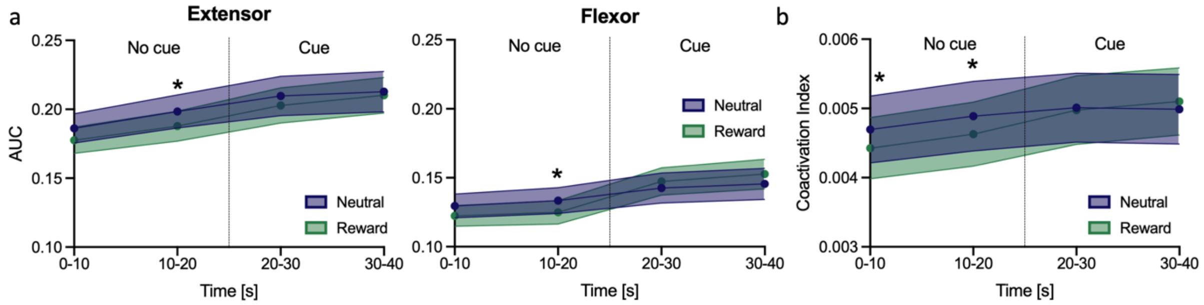


**Supplementary Figure 1. EMG order effects.** Binned AUC of ECR and FCR (a), and the coactivation index (b) of the two muscles. These plots represent the preregistered analysis and are presented here because we implemented an additional normalization (reported in the main text) that was not included in our preregistration due to the discovery of an order effect. Complete statistical results are provided in Supplementary Table 5. Despite the task structure being identical across conditions prior to cue onset, significant differences emerged between neutral and reward conditions in the first two time bins (0-10s, 10-20s). Further analysis suggested that these early differences might reflect an order effect, as transitions between different conditions (neutral-to-reward: 41%; reward-to-neutral: 35%) were more frequent than repetitions of the same condition (neutral-to-neutral: 8%; reward-to-reward: 16%). To test this effect statistically, we examined whether changes in the muscle activity of the first-bin (0–10s) across consecutive trials varied systematically by transition type. For each consecutive trial pair, we computed the difference in first-bin AUC (or coactivation index) between the current and preceding trial and classified each pair by its transition type (neutral-to-neutral, neutral-to-reward, reward-to-neutral, reward-to-reward). These difference scores were submitted to a linear model with transition type as a fixed factor. The AUC of both muscles and their coactivation index were significantly influenced by transition type (ECR: $F(3) = 4.193$, $p = 0.008$, $\eta^2 = 0.12$, 95% CI for $\eta^2$: [0.02, 1.00]; FCR: $F(3) = 3.191$, $p = 0.027$, $\eta^2 = 0.09$, 95% CI for $\eta^2$: [0.01, 1.00]; CI: $F(3) = 5.458$, $p = 0.002$, $\eta^2 = 0.15$, 95% CI for $\eta^2$: [0.04, 1.00]). Changes in early muscle activity differed significantly between control-to-reward and reward-to-control transitions (ECR: $t(94) = -3.286$, $p = 0.008$; FCR: $t(94) = -3.974$, $p = 0.019$; CI: $t(94) = -3.373$, $p = 0.006$). This suggests that the breaks after reward trials were insufficient for full recovery. The unbalanced condition transition frequencies thus offer a plausible explanation for the early between-condition differences observed in panel a. The additional pre-processing step was necessary to account for this order effect in our main analysis. Shaded areas represent standard error of the mean (SEM). Dashed vertical lines indicate cue onset after the first half of tapping (20s). * $p < .05$.

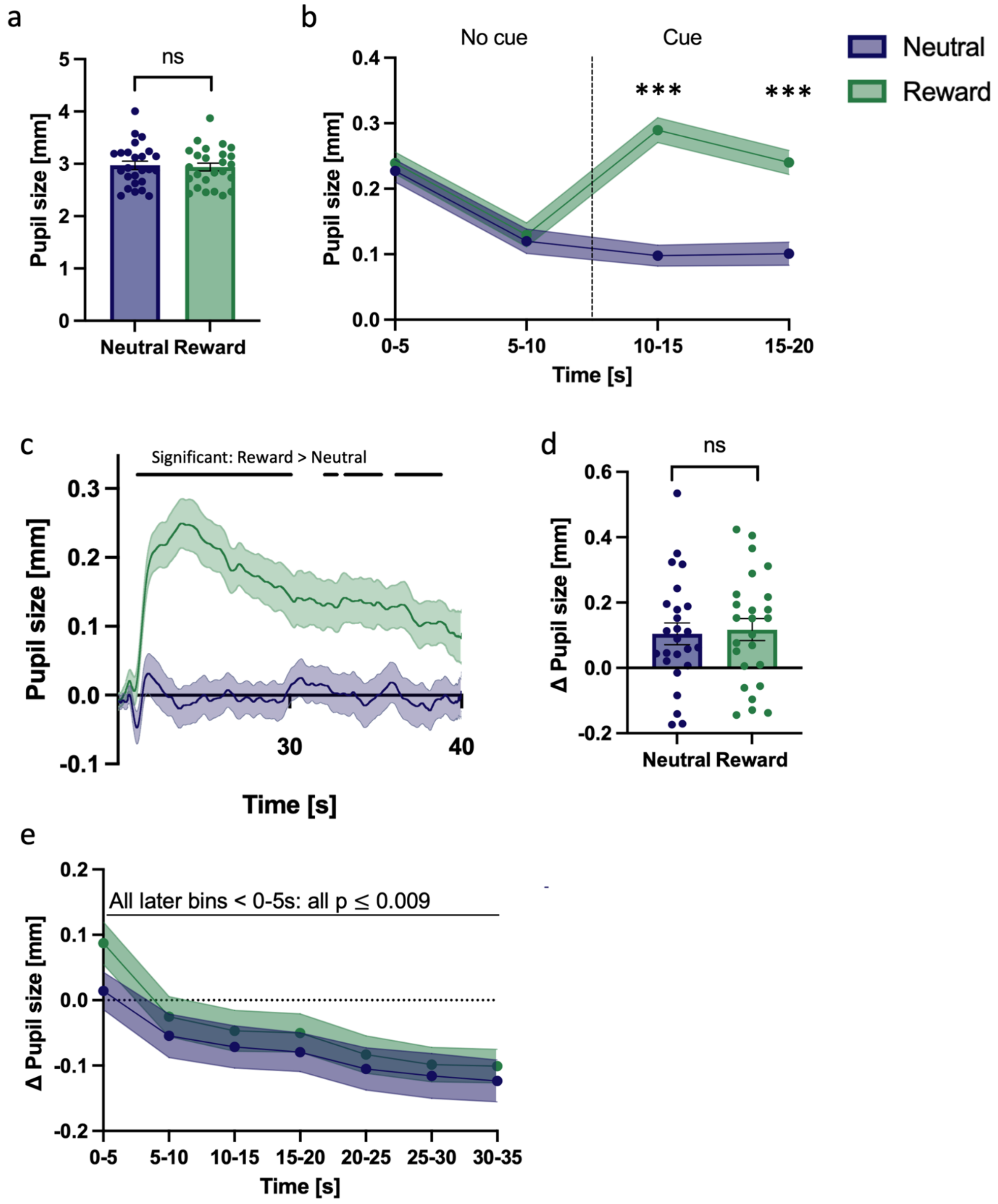


**Supplementary Figure 2. Pupil size across experimental conditions**. (a) Absolute pupil diameter (uncorrected) averaged over the last second of the baseline phase. No significant difference was observed between the neutral and reward conditions (two-tailed paired t-test: $t(24) = 1.567$, $p = 0.130$). (b) Time-binned, baseline-corrected pupil diameter averaged across participants. These binned data was used for the statistical analysis reported in the main text (see Results: Pupil size increases in response to the reward cue). (c) Cue-evoked pupil dilation responses, corrected to the mean pupil size 2.5 s prior to cue onset. Dilation responses were significantly higher in the reward compared to the neutral condition (SPM1D: $z^* = 3.626$; largest cluster $p = 0.013$; smallest cluster $p = 0$). (d) Mean baseline-corrected pupil diameter during the 2.5 s preceding cue onset did not differ significantly between conditions ($t(24) = -0.452$, $p = 0.656$), indicating that the group difference in cue-evoked dilation (panel c) is not attributable to baseline discrepancies. (e) Binned baseline-corrected pupil diameter during the break averaged across participants (see Results: Pupil size increases in response to the reward cue (see Results: Control analyses). For bar plots (a, d): Each dot represents one participant; error bars indicate SEM; "ns" indicates $p > 0.05$. For time series (b, c, e): Shaded areas indicate SEM; dashed vertical lines mark cue onset (b, c); black lines highlight time clusters showing significant differences between conditions (reward vs. neutral, $p < 0.05$; panel c). In panel (e), the horizontal black line indicates that pupil size was significantly lower for all time bins in comparison to the first time bin of the break (all $p \leq 0.009$). In panel (b), *** indicates $p \leq 0.001$.

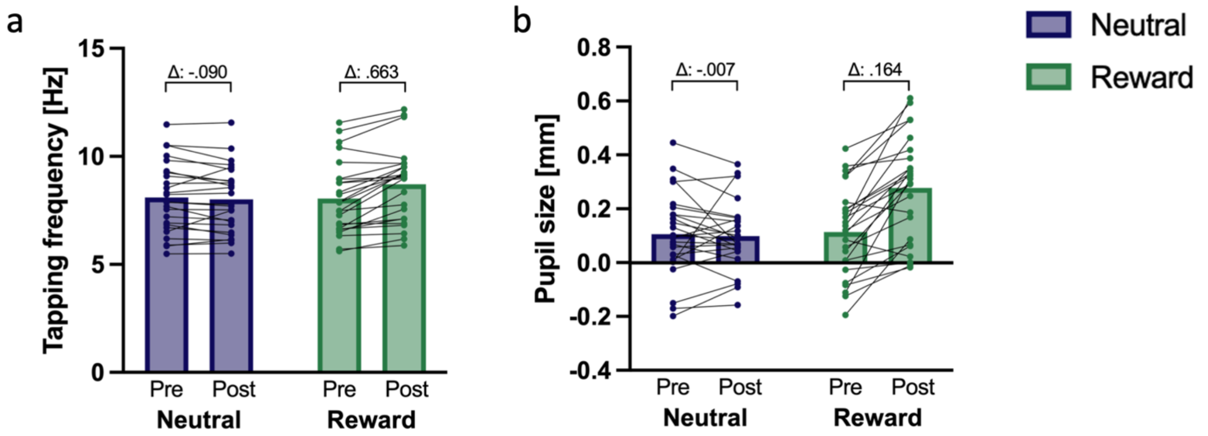


**Supplementary Figure 3. Cue-evoked changes in tapping frequency and pupil size.** Average tapping frequency (a) and pupil size (b) during the 5s before (pre) and the 5s after (post) cue onset, shown separately for the neutral and the reward condition. Each dot represents the average value of one participant; pre and post values of each participant are connected by a black line.

### Supplementary Tables

**Supplementary Table 1. Comparison of model-based and bootstrap confidence intervals for area under the curve of the M. extensor carpi radialis longus (ECR)**. Although the residuals of the LMEM violated normality, (Kolmogorov-Smirnov test with $p < 0.05$), bootstrap validation supported the robustness of our parameter estimates, yielding confidence intervals highly consistent with model-based estimates.

| Parameter | Estimate fixed effects | Model 95% CI | Bootstrap 95% CI |
|---|---|---|---|
| (Intercept) | 1 | [0.952, 1.048] | [0.977, 1.020] |
| 10-20s | 0.063 | [0.014, 0.112] | [0.029, 0.093] |
| 20-30s | 0.120 | [0.071, 0.169] | [0.069, 0.172] |
| 30-40s | 0.143 | [0.094, 0.192] | [0.085, 0.206] |
| Reward | 0.000 | [-0.049, 0.049] | [-0.000, 0.000] |
| 10-20s:Reward | -0.008 | [-0.077, 0.062] | [-0.029, 0.013] |
| 20-30s:Reward | 0.020 | [-0.050, 0.089] | [-0.017, 0.054] |
| 30-40s:Reward | 0.044 | [-0.025, 0.113] | [-0.003, 0.087] |

**Supplementary Table 2. Comparison of model-based and bootstrap confidence intervals for area under the curve of the M. flexor carpi radialis (FCR).** Although the residuals of the LMEM violated normality, (Kolmogorov-Smirnov test with $p < 0.05$), bootstrap validation supported the robustness of our parameter estimates, yielding confidence intervals highly consistent with model-based estimates.

| Parameter | Estimate fixed effects | Model 95% CI | Bootstrap 95% CI |
|---|---|---|---|
| (Intercept) | 1 | [0.947, 1.053] | [0.975, 1.020] |
| 10-20s | 0.029 | [-0.025, 0.083] | [-0.003, 0.063] |
| 20-30s | 0.101 | [0.047, 0.154] | [0.048, 0.160] |
| 30-40s | 0.126 | [0.072, 0.180] | [0.057, 0.204] |
| Reward | -0.000 | [-0.054, 0.054] | [-0.000, 0.000] |
| 10-20s:Reward | -0.008 | [-0.084, 0.068] | [-0.032, 0.016] |
| 20-30s:Reward | 0.111 | [0.035, 0.187] | [0.069, 0.153] |
| 30-40s:Reward | 0.128 | [0.052, 0.205] | [0.073, 0.185] |

**Supplementary Table 3. Comparison of model-based and bootstrap confidence intervals for the coactivation index of the M. flexor carpi radialis and M. extensor carpi radialis longus**. Although the residuals of the LMEM violated normality, (Kolmogorov-Smirnov test with $p < 0.05$), bootstrap validation supported the robustness of our parameter estimates, yielding confidence intervals highly consistent with model-based estimates besides for time bin 10-20s.

| Parameter | Estimate fixed effects | Model 95% CI | Bootstrap 95% CI |
|---|---|---|---|
| (Intercept) | 1 | [0.961, 1.039] | [0.983, 1.010] |
| 10-20s | 0.041 | [-0.002, 0.084] | [0.007, 0.075] |
| 20-30s | 0.078 | [0.035, 0.122] | [0.042, 0.117] |
| 30-40s | 0.079 | [0.035, 0.122] | [0.036, 0.130] |
| Reward | 0.000 | [-0.046, 0.046] | [-0.011, 0.013] |
| 10-20s:Reward | -0.009 | [-0.052, 0.070] | [-0.018, 0.035] |
| 20-30s:Reward | 0.054 | [-0.007, 0.115] | [-0.004, 0.099] |
| 30-40s:Reward | 0.092 | [0.031, 0.153] | [0.043, 0.141] |

**Supplementary Table 4**. **Comparison of model-based and bootstrap confidence intervals for the pupil data analysis during the break.** Since the residuals of the LMEM violated the assumption of normality, we compared model-based and bootstrap confidence interval (see Methods Statistical Analysis for details). Both approaches converged on the same main effects: pupil size decreased significantly over time, and pupil size was larger during the reward condition compared with the first time bin of the neutral condition. However, the confidence intervals of the Time x Condition interaction differed between the two methods (marked in bold). The bootstrap confidence intervals for the interaction terms consistently excluded zero, suggesting a potential diminishing reward effect over time during the break. Given this discrepancy between methods and the exploratory nature of this analysis, these findings should be interpreted with caution.

| Parameter | Estimate fixed effects | Model 95% CI | Bootstrap 95% CI |
|---|---|---|---|
| (Intercept) | 0.014 | [-0.049, 0.078] | [-0.033, 0.060] |
| 5-10s | -0.069 | [-0.107, -0.031] | [-0.107, -0.028] |
| 10-15s | -0.086 | [-0.124, -0.048] | [-0.133, -0.039] |
| 15-20s | -0.094 | [-0.132, -0.055] | [-0.132, -0.053] |
| 20-25s | -0.120 | [-0.158, -0.081] | [-0.165, -0.072] |
| 25-30s | -0.130 | [-0.169, -0.092] | [-0.172, -0.087] |
| 30-35s | -0.138 | [-0.176, -0.099] | [-0.178, -0.096] |
| Reward | 0.073 | [0.019, 0.127] | [0.032, 0.112] |
| 5-10s:reward | -0.043 | **[-0.098, 0.011]** | **[-0.074, -0.013]** |
| 10-15s:reward | -0.048 | **[-0.102, 0.006]** | **[-0.081, -0.016]** |
| 15-20s:reward | -0.044 | **[-0.098, 0.010]** | **[-0.084, -0.007]** |
| 20-25s:reward | -0.051 | **[-0.105, 0.003]** | **[-0.092, -0.007]** |
| 25-30s:reward | -0.055 | **[-0.109, -0.001]** | **[-0.095, -0.012]** |
| 30-35s:reward | -0.050 | **[-0.104, 0.004]** | **[-0.085, -0.012]** |

***Supplementary Table 5. Statistical results of the EMG analysis as preregistered (not normalized to the first time bin).*** *This table presents all statistical analyses from the preregistered approach, organized by EMG measure. For each measure, main effects and interactions from Linear Mixed Effects Models (LMEM) are presented first, followed by post-hoc pairwise comparisons (for the post hoc analysis: only significant results are reported here). The "Factor Level" column indicates either the condition (Neutral/Reward) for time comparisons or the time bin (0-10s, 10-20s, 20-30s, 30-40s) for condition comparisons. Main effects report F-values with degrees of freedom, p-values, partial eta-squared ($\eta_p^2$) effect sizes, and 95% confidence intervals for the $\eta_p^2$. Post-hoc comparisons report parameter estimates, standard errors (SE), t-values, degrees of freedom (df), and p-values. AUC = Area Under the Curve; ECR = Extensor Carpi Radialis; FCR = Flexor Carpi Radialis.*

| **AUC of the ECR - Main effects and interactions of the LMEM** | | | | | | |
|---|---|---|---|---|---|---|
| **Measure** | **Effect** | **F-value** | **df** | **p-value** | $\eta_p^2$ | **95% CI** |
| **ECR AUC** | Time | 25.942 | (3, 168) | < 0.0001 | 0.32 | [0.22, 1.00] |
| | Condition | 7.368 | (1, 168) | 0.007 | 0.04 | [0.01, 1.00] |
| | Time × Condition | 0.417 | (3, 168) | 0.741 | 0.007 | [0.00, 1.00] |
| **AUC of the ECR - Post hoc parwise comparisons:** | | | | | | |
| **Factor level** | **Comparison** | **Estimate** | **SE** | **t-value** | **df** | **p-value** |
| Neutral | 0–10s vs. 10–20s | -0.012 | 0.005 | -2.333 | 168 | .125 |

| | | | | | | |
|---|---|---|---|---|---|---|
| Neutral | 0–10s vs. 20–30s | -0.024 | 0.005 | -4.478 | 168 | .0001 |
| Neutral | 0–10s vs. 30–40s | -0.027 | 0.005 | -5.087 | 168 | < .0001 |
| Neutral | 10–20s vs. 20–30s | -0.011 | 0.005 | -2.145 | 168 | .200 |
| Neutral | 10–20s vs. 30–40s | -0.014 | 0.005 | -2.753 | 168 | .039 |
| Neutral | 20–30s vs. 30–40s | -0.003 | 0.005 | -0.608 | 168 | 1.000 |
| Reward | 0–10s vs. 10–20s | -0.010 | 0.005 | -1.959 | 168 | .311 |
| Reward | 0–10s vs. 20–30s | -0.025 | 0.005 | -4.820 | 168 | <.0001 |
| Reward | 0–10s vs. 30–40s | -0.033 | 0.005 | -6.228 | 168 | < .0001 |
| Reward | 10–20s vs. 20–30s | -0.015 | 0.005 | -2.861 | 168 | .029 |
| Reward | 10–20s vs. 30–40s | -0.022 | 0.005 | -4.268 | 168 | .0002 |
| Reward | 20–30s vs. 30–40s | -0.007 | 0.005 | -1.408 | 168 | .966 |
| 0–10s | Neutral vs. Reward | 0.009 | 0.005 | 1.634 | 168 | .104 |
| 10–20s | Neutral vs. Reward | 0.011 | 0.005 | 2.008 | 168 | .046 |
| 20–30s | Neutral vs. Reward | 0.007 | 0.005 | 1.293 | 168 | .198 |
| 30–40s | Neutral vs. Reward | 0.003 | 0.005 | 0.493 | 168 | .623 |
| **AUC of the FCR- Main effects and interactions of the LMEM:** | | | | | | |
| **Measure** | **Effect** | **F-value** | **df** | **p-value** | $\eta_p^2$ | **95% CI** |
| **FCR AUC** | Time | 36.005 | (3, 168) | < 0.0001 | 0.39 | [0.29, 1.00] |
| | Condition | 0.2735 | (1, 168) | 0.602 | 0.002 | [0.00, 1.00] |
| | Time × Condition | 4.4362 | (3, 168) | 0.005 | 0.07 | [0.01, 1.00] |

| **AUC of the FCR - Post hoc parwise comparisons:** | | | | | | |
|---|---|---|---|---|---|---|
| **Factor level** | **Comparison** | **Estimate** | **SE** | **t-value** | **df** | **p-value** |
| Neutral | 0–10s vs. 10–20s | -0.004 | 0.004 | -1.038 | 168 | 1.00 |
| Neutral | 0–10s vs. 20–30s | -0.013 | 0.004 | -3.415 | 168 | .005 |
| Neutral | 0–10s vs. 30–40s | -0.016 | 0.004 | -4.210 | 168 | .0002 |
| Neutral | 10–20s vs. 20–30s | -0.009 | 0.004 | -2.377 | 168 | .111 |
| Neutral | 10–20s vs. 30–40s | -0.012 | 0.004 | -3.172 | 168 | .011 |
| Neutral | 20–30s vs. 30–40s | -0.003 | 0.004 | -0.795 | 168 | 1.000 |
| Reward | 0–10s vs. 10–20s | -0.003 | 0.004 | -0.698 | 168 | 1.000 |
| Reward | 0–10s vs. 20–30s | -0.025 | 0.004 | -6.547 | 168 | < .0001 |
| Reward | 0–10s vs. 30–40s | -0.030 | 0.004 | -7.966 | 168 | < .0001 |
| Reward | 10–20s vs. 20–30s | -0.022 | 0.004 | -5.848 | 168 | < .0001 |
| Reward | 10–20s vs. 30–40s | -0.0278 | 0.004 | -7.267 | 168 | < .0001 |
| Reward | 20–30s vs. 30–40s | -0.005 | 0.004 | -1.419 | 168 | .947 |
| 0–10s | Neutral vs. Reward | 0.007 | 0.004 | 1.898 | 168 | .059 |
| 10–20s | Neutral vs. Reward | 0.009 | 0.004 | 2.238 | 168 | .027 |
| 20–30s | Neutral vs. Reward | -0.005 | 0.004 | -1.233 | 168 | .219 |
| 30–40s | Neutral vs. Reward | -0.007 | 0.004 | -1.857 | 168 | .065 |
| **Coactivation Index of the ECR and the FCR - Main effects and interactions of the LMEM** | | | | | | |
| **Measure** | **Effect** | **F-value** | **df** | **p-value** | $\eta_p^2$ | **95% CI** |

| | | | | | | |
|---|---|---|---|---|---|---|
| **ECR-FCR Overlap** | Time | 22.270 | (3, 144) | < 0.0001 | 0.32 | [0.21, 1.00] |
| | Condition | 1.954 | (1, 24) | 0.175 | 0.08 | [0.00, 1.00] |
| | Time × Condition | 3.835 | (3, 144) | 0.011 | 0.07 | [0.01, 1.00] |
| **Coactivation Index of the ECR and the FCR - Post hoc pairwise comparisons** | | | | | | |
| **Factor level** | **Comparison** | **Estimate** | **SE** | **t-value** | **df** | **p-value** |
| Neutral | 0–10s vs. 10–20s | -0.0002 | 0.0001 | -2.037 | 144 | .261 |
| Neutral | 0–10s vs. 20–30s | -0.0003 | 0.0001 | -3.327 | 144 | .006 |
| Neutral | 0–10s vs. 30–40s | -0.0003 | 0.0001 | -3.087 | 144 | .015 |
| Neutral | 10–20s vs. 20–30s | -0.0001 | 0.0001 | -1.290 | 144 | 1.000 |
| Neutral | 10–20s vs. 30–40s | -0.0001 | 0.0001 | -1.050 | 144 | 1.000 |
| Neutral | 20–30s vs. 30–40s | 0.0000 | 0.0001 | 0.240 | 144 | 1.000 |
| Reward | 0–10s vs. 10–20s | -0.0002 | 0.0001 | -2.138 | 144 | .205 |
| Reward | 0–10s vs. 20–30s | -0.0006 | 0.0001 | -5.803 | 144 | < .0001 |
| Reward | 0–10s vs. 30–40s | -0.0007 | 0.0001 | -7.125 | 144 | < .0001 |
| Reward | 10–20s vs. 20–30s | -0.0002 | 0.0001 | -3.664 | 144 | .0002 |
| Reward | 10–20s vs. 30–40s | -0.0005 | 0.0001 | -4.987 | 144 | <.0001 |
| Reward | 20–30s vs. 30–40s | -0.0001 | 0.0001 | -1.323 | 144 | 1.000 |
| 0–10s | Neutral vs. Reward | 0.0002 | 0.0001 | 2.341 | 84 | .022 |
| 10–20s | Neutral vs. Reward | 0.0003 | 0.0001 | 2.258 | 84 | .027 |

| | | | | | | |
|---|---|---|---|---|---|---|
| 20–30s | Neutral vs. Reward | 0.0000 | 0.0001 | 0.306 | 84 | .761 |
| 30–40s | Neutral vs. Reward | -0.0001 | 0.0001 | -0.979 | 84 | .330 |
